# An Aging Risk-Factor Scale: Biomarkers of Renal Disease and Anemia are Primary Predictors of Three-Year Survival in Common Marmosets (*Callithrix jacchus*)

**DOI:** 10.64898/2026.08.06.743332

**Authors:** Juan Pablo Arroyo, Aaryn C. Mustoe, Kelly R. Reveles, Kathleen M. Brasky, Donna Perry, Lidia Cervantes, Addaline Alvarez, Clarissa Hinojosa, Jessica Greig, Alexana J. Hickmott, Benjamin J. Ridenhour, Katherine R. Amato, Michael L. Power, Corinna N. Ross

## Abstract

Valid animal models are needed to evaluate how age-related changes in kidney function influence healthspan. Aging marmosets frequently develop renal insufficiency with anemia and exhibit reductions in body mass and metabolic rate. However, it remains unclear which age-related changes predict survival and which thresholds indicate increased mortality risk. We prospectively evaluated age, body composition, resting energy expenditure, hematology, and blood chemistry as predictors of 3-year survival in female and male marmosets (n = 66), 2–16 years of age. Objectives were to identify prognostic markers, define high-risk thresholds, and to develop and test a composite risk-factor scale for mortality screening in captivity. A 10-variable model showed the best predictive performance in multivariable Cox proportional hazards modeling, and was retained for further analysis (concordance = 0.881, p < 0.001). ROC curves using Youden’s Index and AUC identified high-risk thresholds for predictors in the multivariable model, and threshold-defined categories were evaluated by Kaplan–Meier survival analysis. The 10 binary risk-factors were combined into a composite scale scored from 0 to 10 and tested with Cox regression. The scale explained approximately 42% of variance in survival and each additional risk factor increased mortality risk 1.75-fold (95% CI: 1.43–2.14, p < 0.001). Marmosets with ≥7 risk factors exhibited a 19-month reduction in survival, and this high-risk threshold predicted 3-year survival with 89.4% accuracy. Results support the scale as a screening tool for mortality risk and highlight the high prevalence of age-associated renal disease and anemia in marmosets.

## Introduction

The common marmoset (*Callithrix jacchus*) is a small South American nonhuman primate (NHP) that has seen rapidly increasing demand as a biomedical model organism [1, 2], in part because its close genetic relatedness to humans confers substantial translational relevance [3]. Marmosets weigh approximately 300–450 g, reach reproductive maturity at ∼1.5 years, and typically produce litters of 2–3 offspring at 5–6 month intervals. They have a comparatively short primate life-course, with animals considered aged by 8–12 years [4, 5]. Collectively, these life-history traits offer practical advantages over larger, longer-lived NHP models and have supported the broad adoption of marmosets in neuroscience, aging biology, and infectious disease research [1, 6, 7].

Marmosets exhibit strong potential as a model for studying chronic kidney disease (CKD) and anemia. Experimentally induced renal insufficiency via partial nephrectomy (i.e., 5/6 nephrectomy) in marmosets produces pathologies that closely resemble chronic renal failure in humans, including elevations in serum urea nitrogen (SUN) and creatinine, along with bone marrow hypocellularity and anemia [8]. In natural aging, marmosets show declines in metabolic rate [9] and body mass [10], with increased mortality risk attributable to CKD [11]. Consistent with this clinical pattern, aged marmosets frequently exhibit renal structural lesions characteristic of renal aging such as glomerulosclerosis, interstitial fibrosis, and arteriolosclerosis [12, 13]. Together, these features support the marmoset as a translational platform for mechanistic and interventional geroscience studies (e.g., rapamycin, GLP-1 receptor agonists) aimed at understanding and modifying aging-related CKD and associated anemia, and obesity [6, 14].

Across mammals, aging is accompanied by clinically silent, yet progressive loss of nephrons and a decline in renal functional reserve (e.g., decreased capacity in glomerular filtration rate) leading to CKD [15–17]. Because the kidneys are essential for sodium and water regulation, pH (i.e., acid-base balance), and endocrine support of erythropoiesis (i.e., Norn cell production of erythropoietin); renal impairment can alter fluid status (i.e., balance of body water) in body composition, and have downstream effects that are highly relevant to late-life vulnerability. Decline in renal functional reserve promotes fluid volume dysregulation (i.e., homeostatic failure at maintaining fluid concentration balance), leading to fluid retention (i.e., extracellular accumulation of fluid) and overload (i.e., state of fluid exceeding physiological thresholds); resulting in adverse clinical outcomes [18, 19]. In parallel, CKD is a major driver of anemia, through reduced renal erythropoietin production with additional contributions from disordered iron handling and impaired erythropoietic responsiveness [20, 21]. Aging-associated, chronic inflammation (“inflammaging”) can accelerate chronic disease progression and disrupt normal regulation of hematopoiesis. As a result, inflammation, kidney dysfunction, altered fluid status, and anemia often cluster with age [22–24].

We previously reported reference ranges for blood chemistry and hematology parameters in common marmosets. Age-associated shifts were documented on otherwise healthy older animals in hepatic, pancreatic, renal, and erythrocyte-related markers, highlighting increased susceptibility to kidney disease and anemia with age [25]. However, it has remained unclear whether these age-related changes prospectively predict survival, and which values should be interpreted as clinically meaningful thresholds of elevated mortality risk. This uncertainty limits both research design and clinical decision making in captive marmoset colonies, underscoring the need for a validated, marmoset-specific risk scale that can be implemented as a practical screening tool.

A screening scale could help identify animals with subclinical dysregulation or early-stage disease states that may be more responsive to intervention than overt, late-stage illness. In colony practice, a risk scale could prompt enhanced monitoring and targeted diagnostic follow-up, guide decisions about exclusion from experimental protocols when imminent death would confound study endpoints, and support timely enrollment into welfare or palliative care pathways. For geroscience research, improved resolution of mortality risk across key physiological domains would enable more precise baseline stratification, covariate adjustment for “physiological age”, and selective enrollment into intervention trials based on predefined risk strata.

Accordingly, in the present study, age, body composition, resting metabolic rate, blood chemistry, and hematology variables were measured at baseline and evaluated prospectively as predictors of 3-year survival in common marmosets. Our objectives were to: 1) identify informative prognostic markers, 2) define empirically derived high-risk thresholds, and 3) develop and test a composite risk-factor scale as a screening tool for mortality risk in captivity.

## Materials and Methods

### Subjects

This study examined 64 adult common marmosets (*Callithrix jacchus*): 33 females and 33 males, spanning 2–16 years of age. All animals were housed at the Southwest National Primate Research Center (SNPRC), Texas Biomedical Research Institute (an AAALAC-accredited institution), under standardized barrier husbandry conditions [26, 27]. Animals were provided two base diets daily, the Harlan Teklad purified marmoset diet and the Mazuri marmoset diet [28]. Daily dietary enrichment was provided in small quantities using a rotating selection of items (e.g., dried cranberries and legumes), and water was available ad libitum.

The marmosets were housed as male–female pairs, and no females were gravid or postpartum during the study period. None of the animals had participated in prior studies that could have an effect on markers and outcomes measured in this investigation. The research protocols were approved by the SNPRC animal care and use committee (IACUC #1743 CJ), and complied with all applicable U.S. laws regarding animal research.

### Body Composition and Metabolic Rates

Whole-body composition was assessed to quantify lean mass, fat mass, and fluid status by placing unsedated animals in an EchoMRI quantitative magnetic resonance (QMR) system (Houston, TX, USA) [29] and performing a 2-minute scan [10]. Resting energy expenditure (REE) was measured using a Sable Systems FoxBox (Las Vegas, NV, USA) flow-through respirometry system. The marmosets fasted for 2 hours and then were placed in a metabolic chamber maintained at 28.9°C with a controlled airflow of 1,000 mL/min for a 2-hour recording period. REE (kcal/day) was calculated from VO₂ and VCO₂ [30] using a modified Weir equation [31]: REE = (5.46 × VO₂ + 1.75 × VCO₂) / 60.

### Hematology and Blood chemistry

Blood was collected once at baseline from each marmoset during a morning physical examination, as part of routine colony health monitoring. Animals were fasted at 8:00 am and sedated with intramuscular ketamine (20 mg/kg) before veterinary assessment. For complete blood count (CBC) and serum chemistry analyses, 1.5 mL of blood was drawn from the femoral vein; 0.5 mL was placed in an EDTA tube for CBC, and 1.0 mL was placed in a serum separator tube for chemistry. Samples were transported to the SNPRC Veterinary Clinical Pathology Core, where CBCs were analyzed with a UniCel DxH 800 Coulter Cellular Analysis System and serum chemistry panels were analyzed with a DxC 700 AU Chemistry Analyzer.

### Statistical Analysis

The first objective was to prospectively evaluate health markers as predictors of 3-year survival. To address this, survival analyses were conducted for body composition, resting metabolic rate, hematology, and blood chemistry variables, with separate Cox proportional hazards models for each continuous marker. These univariable Cox models served as an initial step to assess associations across the full distribution of each variable. Subsequently, multiple numbers and combinations of variables were evaluated in multivariable Cox proportional hazards models.

Because many predictors were biologically and statistically correlated, a multivariable model that included a large number of predictors was not feasible without introducing substantial multicollinearity and unstable coefficient estimates. In addition, the number of deaths within 3 years was modest relative to the total number of candidate predictors, increasing the risk of overfitting and optimistic performance if the model were over-parameterized. Therefore, multivariable modeling was intentionally constrained to a parsimonious set of variables selected to represent the major physiological domains of age, body composition, kidney disease, anemia, and metabolic rate, rather than including multiple partially redundant measures from the same domain. Multiple combinations and numbers of variables were evaluated. Within these constraints, a 10-marker model provided the strongest overall predictive performance in the present dataset and was carried forward for additional analyses.

Lack of significance for a continuous variable was not interpreted as absence of prognostic value, as standard Cox models assume a linear relationship on the log-hazard scale [32]; whereas some biomarkers may exhibit nonlinear or threshold-dependent relationships with mortality, such that elevated risk becomes apparent only beyond a specific cut-point [33]. Accordingly, the second objective was to define risk-categorization thresholds for variables included in the final multivariable model. Receiver operating characteristic (ROC) curve analysis, the Youden index, and area under the curve (AUC) were used to identify cut-points with the highest discriminatory performance. Based on these ROC-derived thresholds, each variable included in the multivariable model was dichotomized into binary risk categories (low vs. high risk) and evaluated with Kaplan–Meier analysis using the log-rank, Gehan, Tarone–Ware, and Peto–Peto tests.

The third objective was to develop and evaluate a composite risk-factor scale for identifying individuals at risk of death within 3 years. After ROC-based dichotomization and Kaplan–Meier evaluation of threshold prognostic utility, the 10 variables in the final multivariable Cox model were incorporated into a composite scale. Because the primary aim was risk stratification and screening rather than estimation of the independent causal effect of each marker, the scale was constructed as a simple binary deficit count spanning major physiological domains [34]. This 10-item risk scale was scored by summing the presence (1) or absence (0) of each risk factor (total score range: 0–10). The scale was then tested as a predictor of survival using Cox regression. ROC analysis was subsequently used to define three risk strata by number of risk factors present: low risk (0–1), medium risk (2–6), and high risk (≥7). Its performance as a screening tool for 3-year survival was further evaluated with metrics of accuracy, sensitivity, specificity, positive predictive value, and negative predictive value [35, 36]. Analyses were performed in R version 4.1 [37] through jamovi version 2.3.17 [38], including the epiR package [39] and the ClinicoPath jamovi module [40].

After associations among variables were identified through regression analyses, path analysis was used as an extension of regression to test alternative models of intervariable relationships. This approach enabled evaluation of directional effects for individual paths within each model [41]. Path models were estimated in jamovi PATHj module [42] using the lavaan [43] and semPlot [44] R packages. An overall path model was produced to examine and summarize how the principal survival predictors across the most salient health domains in the dataset were related to one another. Model fit was evaluated using the Chi-square goodness of fit test, root mean square error of approximation (RMSEA), comparative fit index (CFI), Tucker–Lewis index (TLI), and standardized root mean square residual (SRMR) [45].

## Results

Females exhibited higher fat mass (females: *M* = 26.6, SD = 17.74; males: *M* = 18.33, SD = 14.4; *F* = 4.19, p=0.045), and higher fat percentage (females: *M* = 5.84, SD = 3.25; males: *M* = 4.22, SD = 3.04; *F* = 4.22, p=0.044) than males. There were no other sex differences in body composition, metabolic rate, blood chemistry or hematology parameters. Therefore, female and male data were analyzed together.

At the 3-year follow-up, 48 animals remained alive and 18 did not survive. Of the 18 non-survivors, 12 were euthanized based on veterinary discretion. Necropsy findings indicated that kidney disease was present in 15 of the 18 non-survivors (83.3%). Of these 15 cases, 12 deaths, including euthanasia, were attributed to kidney disease complications (80.0%). Kidney disease at necropsy was associated with increased odds of concurrent anemia: 14 of 15 animals with confirmed kidney disease (93.3%) also had anemia at necropsy (OR=28.0, 95% CI: 1.21–648.81, p=0.038; McFadden R² = 0.312). Among the 12 animals with deaths attributed to kidney disease complications, 11 (91.7%) also had anemia confirmed at necropsy.

There were no significant differences in age (*F* = 1.31, p=0.257), body weight (*F* = 1.26, p=0.265), lean mass (*F* = 0.790, p=0.378), or fat mass (*F* = 3.39, p=0.070) between non-survivors and survivors. Non-survivors had lower REE (non-survivors: *M* = 26.94, SD = 7.35; survivors: *M* = 32.74, SD = 10.24; *F* = 4.59, p=0.036), lower fat percentage (non-survivors: *M* = 3.66, SD = 2.56; survivors: *M* = 5.48, SD = 3.32; *F* = 4.01, p=0.05), and higher fluid retention (non-survivors: *M* = 4.08, SD = 1.41; survivors: *M* = 3.23, SD = 1.27; *F* = 5.07, p=0.028) than survivors (Table 1).

**Table 1.** Age, body composition, and resting energy expenditure by 3-year survival status.

| Parameter | Unit | Non-survivors<br>(n); M ± SD | Survivors<br>(n); M ± SD | F statistic | df | p-value |
| --- | --- | --- | --- | --- | --- | --- |
| **Chronological age** |  |  |  |  |  |  |
| Age | years | (n = 17); 8.76 ± 3.63 | (n = 48); 7.75 ± 2.96 | 1.31 | 1, 63 | 0.257 |
| **Body composition** |  |  |  |  |  |  |
| Body weight | g | (n = 16); 410.94 ± 68.74 | (n = 48); 429.81 ± 54.38 | 1.26 | 1, 62 | 0.265 |
| Lean mass | g | (n = 16); 355.56 ± 52.55 | (n = 48); 366.31 ± 37.85 | 0.790 | 1, 62 | 0.378 |
| Fat mass | g | (n = 16); 15.98 ± 12.89 | (n = 48); 24.63 ± 17.19 | 3.39 | 1, 62 | 0.070 |
| Fat percentage | % | (n = 16); 3.66 ± 2.56 | (n = 48); 5.48 ± 3.32 | 4.01 | 1, 62 | 0.050* |
| Fluid retention | g | (n = 16); 4.08 ± 1.41 | (n = 48); 3.23 ± 1.27 | 5.07 | 1, 62 | 0.028* |
| **Metabolic rate** |  |  |  |  |  |  |
| Resting energy expenditure (REE) | kcal/day | (n = 17); 26.94 ± 7.35 | (n = 48); 32.74 ± 10.24 | 4.59 | 1, 63 | 0.036* |

Values are (n); mean ± SD. p-values are from one-way ANOVA comparing baseline values between non-survivors and survivors at 3-year follow-up.
\*statistically significant

In blood chemistry, non-survivors had higher serum urea nitrogen (SUN) (non-survivors: *M* = 61.6, SD = 59.36; survivors: *M* = 20.77, SD = 5.41; *F* = 22.4, p<0.001), higher creatinine (non-survivors: *M* = 0.88, SD = 0.80; survivors: *M* = 0.39, SD = 0.12; *F* = 16.6, p<0.001), higher SUN/creatinine ratio (non-survivors: *M* = 71.67, SD = 38.15; survivors: *M* = 56.11, SD = 18.46; *F* = 4.46, p=0.039), higher calcium (non-survivors: *M* = 10.0, SD = 0.784; survivors: *M* = 9.57, SD = 0.666; *F* = 4.43, p=0.039), lower chloride (non-survivors: *M* = 102.8, SD = 3.93; survivors: *M* = 105.06, SD = 3.25; *F* = 4.99, p=0.029), higher phosphorus (non-survivors: *M* = 4.13, SD = 1.22; survivors: *M* = 3.35, SD = 0.81; *F* = 8.26, p=0.006), and higher potassium (non-survivors: *M* = 3.63, SD = 0.68; survivors: *M* = 3.29, SD = 0.42; *F* = 5.25, p=0.026) than survivors (Table 2).

**Table 2.** Blood chemistry parameters by 3-year survival status.

| Parameter | Unit | Non-survivors (n = 15)<br>M ± SD | Survivors (n = 47)<br>M ± SD | F statistic | df | p-value |
| --- | --- | --- | --- | --- | --- | --- |
| **Metabolic analytes** |  |  |  |  |  |  |
| Glucose | mg/dL | 151.1±50.5 | 153.23±46.56 | 0.024 | 1, 60 | 0.878 |
| Cholesterol | mg/dL | 183.8±60.83 | 172.26±113.89 | 0.140 | 1, 60 | 0.709 |
| **Renal function** |  |  |  |  |  |  |
| Serum urea nitrogen (SUN) | mg/dL | 61.6 ± 59.36 | 20.77 ± 5.41 | 22.4 | 1, 60 | <0.001* |
| Creatinine | mg/dL | 0.88 ± 0.80 | 0.39 ± 0.12 | 16.6 | 1, 60 | <0.001* |
| SUN/creatinine ratio |  | 71.67 ± 38.15 | 56.11 ± 18.46 | 4.46 | 1, 60 | 0.039* |
| **Electrolytes and acid-base markers** |  |  |  |  |  |  |
| Sodium | meq/L | 149.4±3.25 | 150.02±2.82 | 0.514 | 1, 60 | 0.476 |
| Potassium | meq/L | 3.63 ± 0.68 | 3.29 ± 0.42 | 5.25 | 1, 60 | 0.026* |
| Chloride | meq/L | 102.8 ± 3.93 | 105.06 ± 3.25 | 4.99 | 1, 60 | 0.029* |
| Carbon dioxide | meq/L | 24.8±6.34 | 24±4.78 | 0.271 | 1, 60 | 0.605 |
| Anion gap | meq/L | 25.43±6.53 | 24.25±7.42 | 0.302 | 1, 60 | 0.585 |
| **Minerals** |  |  |  |  |  |  |
| Calcium | mg/dL | 10.0 ± 0.784 | 9.57 ± 0.666 | 4.43 | 1, 60 | 0.039* |
| Phosphorus | mg/dL | 4.13 ± 1.22 | 3.35 ± 0.81 | 8.26 | 1, 60 | 0.006* |
| **Protein profile** |  |  |  |  |  |  |
| Total protein | g/dL | 6.11±0.65 | 6.27±0.68 | 0.634 | 1, 60 | 0.429 |
| Albumin | g/dL | 1.25±0.66 | 1.33±0.56 | 0.034 | 1, 60 | 0.854 |
| Globulin | g/dL | 3.43±1.85 | 3.16±1.54 | 0.300 | 1, 60 | 0.586 |
| Albumin/globulin ratio |  | 1.25±0.66 | 1.33±0.56 | 0.237 | 1, 60 | 0.628 |
| **Hepatobiliary enzymes and pigments** |  |  |  |  |  |  |
| Total bilirubin | mg/dL | 0.14±0.05 | 0.12±0.04 | 2.30 | 1, 60 | 0.134 |
| Alkaline phosphatase | IU/L | 110.07±88.87 | 78.92±41.02 | 3.52 | 1, 60 | 0.065 |
| Alanine aminotransferase | U/L | 58.67±92.04 | 30.92±80.36 | 1.26 | 1, 60 | 0.265 |
| Aspartate aminotransferase | U/L | 166±66.05 | 142.6±86.33 | 0.925 | 1, 60 | 0.340 |

Values are mean ± SD. p-values are from one-way ANOVA comparing baseline values between non-survivors and survivors at 3-year follow-up. \*statistically significant

In hematology, non-survivors had higher eosinophils (non-survivors: *M* = 0.13, SD = 0.16; survivors: *M* = 0.05, SD = 0.08; *F* = 6.37, p=0.014), higher eosinophils percentage (non-survivors: *M* = 2.92, SD = 2.25; survivors: *M* = 1.33, SD = 1.26; *F* = 12.6, p<0.001), lower hematocrit (non-survivors: *M* = 34.06, SD = 11.47; survivors: *M* = 45.26, SD = 5.75; *F* = 26.5, p<0.001), lower hemoglobin (non-survivors: *M* = 10.88, SD = 3.68; survivors: *M* = 14.30, SD = 1.80; *F* = 24.6, p<0.001), and lower RBC (non-survivors: *M* = 5.02, SD = 1.60; survivors: *M* = 6.80, SD = 0.90; *F* = 30.7, p<0.001) than survivors (Table 3).

**Table 3.** Hematology parameters by 3-year survival status.

| Parameter | Unit | Non-survivors (n = 16)<br>M ± SD | Survivors (n = 48)<br>M ± SD | F statistic | df | p-value |
| --- | --- | --- | --- | --- | --- | --- |
| **Leukogram** |  |  |  |  |  |  |
| White blood cell count (WBC) | 10<sup>3</sup>/uL | 4.19±1.66 | 4.71±1.53 | 1.33 | 1, 62 | 0.253 |
| Neutrophils | 10<sup>3</sup>/uL | 1.76±0.78 | 1.8±0.91 | 0.033 | 1, 62 | 0.857 |
| Neutrophils (%) | % | 42.99±17 | 37.78±13.04 | 1.57 | 1, 62 | 0.215 |
| Lymphocytes | 10<sup>3</sup>/uL | 2.14±1.14 | 2.66±1.04 | 2.84 | 1, 62 | 0.097 |
| Lymphocytes (%) | % | 50.06±18.23 | 56.99±14.47 | 2.41 | 1, 62 | 0.126 |
| Monocytes | 10<sup>3</sup>/uL | 0.14±0.09 | 0.17±0.14 | 0.536 | 1, 62 | 0.467 |
| Monocytes (%) | % | 3.49±1.76 | 3.42±1.98 | 0.019 | 1, 62 | 0.890 |
| Eosinophils | 10<sup>3</sup>/uL | 0.13 ± 0.16 | 0.05 ± 0.08 | 6.37 | 1, 62 | 0.014* |
| Eosinophils (%) | % | 2.92 ± 2.25 | 1.33 ± 1.26 | 12.6 | 1, 62 | <0.001* |
| Basophils | 10<sup>3</sup>/uL | 0.0063±0.03 | 0.0083±0.03 | 0.070 | 1, 62 | 0.792 |
| Basophils (%) | % | 0.54±0.62 | 0.5±0.34 | 0.141 | 1, 62 | 0.708 |
| **Erythrogram** |  |  |  |  |  |  |
| Red blood cell count (RBC) | 10<sup>6</sup>/uL | 5.02 ± 1.60 | 6.80 ± 0.90 | 30.7 | 1, 62 | <0.001* |
| Hemoglobin | g/dL | 10.88 ± 3.68 | 14.30 ± 1.80 | 24.6 | 1, 62 | <0.001* |
| Hematocrit | % | 34.06 ± 11.47 | 45.26 ± 5.75 | 26.5 | 1, 62 | <0.001* |
| Mean corpuscular volume (MCV) | fL | 67.32±3.71 | 66.74±3.2 | 0.363 | 1, 62 | 0.549 |
| Mean corpuscular hemoglobin (MCH) | pg | 21.48±1.33 | 21.09±1.05 | 1.44 | 1, 62 | 0.234 |
| Mean corpuscular hemoglobin concentration (MCHC) | g/dL | 31.91±0.62 | 31.6±0.72 | 2.39 | 1, 62 | 0.127 |
| Red cell distribution width (RDW) | % | 14.11±1.52 | 14.7±1.25 | 2.40 | 1, 62 | 0.127 |
| Nucleated red blood cells (NRBC, %) | % | 0.37±0.29 | 0.92±2.75 | 0.636 | 1, 62 | 0.428 |
| **Platelet parameters** |  |  |  |  |  |  |
| Platelet count | 10<sup>3</sup>/uL | 639.69±199.22 | 547.17±163.34 | 3.44 | 1, 62 | 0.068 |
| Mean platelet volume (MPV) | fL | 7.89±0.86 | 8.18±0.87 | 1.40 | 1, 62 | 0.241 |

Values are mean ± SD. p-values are from one-way ANOVA comparing baseline values between non-survivors and survivors at 3-year follow-up. \*statistically significant

### Objective 1: Continuous health markers as predictors of 3-year survival

In univariable Cox proportional hazards models, several markers were significantly associated with survival. Lower REE predicted higher mortality risk (HR=0.92, 95% CI 0.85–0.99, p=0.027), as did higher fluid (HR=1.38, 95% CI 1.07–1.78, p=0.014) (Table 4). Among blood chemistry variables, higher SUN (HR = 1.03, 95% CI 1.02–1.05, p < 0.001), creatinine (HR = 9.30, 95% CI 2.92–29.65, p < 0.001), SUN/creatinine ratio (HR = 1.02, 95% CI 1.01–1.04, p = 0.011), alkaline phosphatase (HR = 1.01, 95% CI 1.00–1.02, p = 0.011), total bilirubin (HR = 3.61, 95% CI 1.27–10.25, p = 0.016), phosphorus (HR = 2.09, 95% CI 1.23–3.55, p = 0.006), calcium (HR = 2.37, 95% CI 1.13–5.00, p = 0.023), and potassium (HR = 3.64, 95% CI 1.48–8.96, p = 0.005) were associated with increased mortality risk, whereas higher chloride was associated with lower risk (HR = 0.82, 95% CI 0.70–0.96, p = 0.014) (Table 4). Lower hemoglobin (HR = 0.60, 95% CI 0.49–0.74, p < 0.001) was associated with increased risk. Among hematologic variables, lower hematocrit (HR = 0.85, 95% CI 0.79–0.90, p < 0.001) and lower red blood cell count (RBC) (HR = 0.34, 95% CI 0.22–0.52, p < 0.001) predicted greater mortality risk, while higher platelets (HR = 1.00, 95% CI 1.00–1.01, p = 0.027), eosinophils (HR = 39.97, 95% CI 2.09–763.23, p = 0.014), and eosinophil percentage (HR = 1.47, 95% CI 1.19–1.82, p < 0.001) were also associated with increased risk (Table 4).

**Table 4.** Cox proportional hazards regression for continuous predictors of 3-year survival.

| Parameter | Unit | Hazard ratio (HR) | 95% CI | p-value | Association with mortality |
| --- | --- | --- | --- | --- | --- |
| **Chronological age** |  |  |  |  |  |
| Age | years | 1.13 | 0.96–1.32 | 0.150 | NS |
| **Body composition** |  |  |  |  |  |
| Body weight | g | 0.99 | 0.98–1.00 | 0.198 | NS |
| Lean mass | g | 1.00 | 0.98–1.01 | 0.510 | NS |
| Fat mass | g | 0.96 | 0.93–1.00 | 0.056 | NS |
| Fat percentage | % | 0.89 | 0.75–1.06 | 0.180 | NS |
| Fluid retention | g | 1.38 | 1.07–1.78 | 0.014* | Increased hazard |
| **Metabolic rate** |  |  |  |  |  |
| Resting energy expenditure (REE) | kcal/day | 0.92 | 0.85–0.99 | 0.027* | Reduced hazard |
| **Blood chemistry** |  |  |  |  |  |
| **Metabolic analytes** |  |  |  |  |  |
| Glucose | mg/dL | 1.00 | 0.99–1.01 | 0.972 | NS |
| Cholesterol | mg/dL | 1.00 | 1.00–1.00 | 0.695 | NS |
| **Renal function** |  |  |  |  |  |
| Serum urea nitrogen (SUN) | mg/dL | 1.03 | 1.02–1.05 | <0.001* | Increased hazard |
| Creatinine | mg/dL | 9.30 | 2.92–29.65 | <0.001* | Increased hazard |
| SUN/creatinine ratio |  | 1.02 | 1.01–1.04 | 0.011* | Increased hazard |
| **Electrolytes and acid-base markers** |  |  |  |  |  |
| Sodium | meq/L | 0.93 | 0.79–1.01 | 0.415 | NS |
| Potassium | meq/L | 3.64 | 1.48–8.96 | 0.005* | Increased hazard |
| Chloride | meq/L | 0.82 | 0.70–0.96 | 0.014* | Reduced hazard |
| Carbon dioxide | meq/L | 1.02 | 0.92–1.13 | 0.689 | NS |
| Anion gap | meq/L | 1.02 | 0.96–1.09 | 0.489 | NS |
| **Minerals** |  |  |  |  |  |
| Calcium | mg/dL | 2.37 | 1.13–5.00 | 0.023* | Increased hazard |
| Phosphorus | mg/dL | 2.09 | 1.23–3.55 | 0.006* | Increased hazard |
| **Protein profile** |  |  |  |  |  |
| Total protein | g/dL | 0.76 | 0.36–1.61 | 0.469 | NS |
| Albumin | g/dL | 0.89 | 0.30–2.60 | 0.825 | NS |
| Globulin | g/dL | 1.07 | 0.80–1.43 | 0.636 | NS |
| Albumin/globulin ratio |  | 0.81 | 0.35–1.86 | 0.620 | NS |
| **Hepatobiliary enzymes and pigments** |  |  |  |  |  |
| Total bilirubin | mg/dL | 3.61 | 1.27–10.25 | 0.016* | Increased hazard |
| Alkaline phosphatase | IU/L | 1.01 | 1.00–1.02 | 0.011* | Increased hazard |
| Alanine aminotransferase | U/L | 1.00 | 1.00–1.01 | 0.259 | NS |
| Aspartate aminotransferase | U/L | 1.00 | 1.00–1.01 | 0.370 | NS |
| **Hematology** |  |  |  |  |  |
| **Leukogram** |  |  |  |  |  |
| White blood cell count (WBC) | 10<sup>3</sup>/uL | 0.80 | 0.56–1.13 | 0.207 | NS |
| Neutrophils | 10<sup>3</sup>/uL | 0.95 | 0.54–1.69 | 0.868 | NS |
| Neutrophils (%) | % | 1.03 | 0.99–1.07 | 0.121 | NS |
| Lymphocytes | 10<sup>3</sup>/uL | 0.58 | 0.33–1.02 | 0.059 | NS |
| Lymphocytes (%) | % | 0.97 | 0.94–1.00 | 0.069 | NS |
| Monocytes | 10<sup>3</sup>/uL | 0.20 | 0.00–15.48 | 0.470 | NS |
| Monocytes (%) | % | 1.03 | 0.80–1.33 | 0.825 | NS |
| Eosinophils | 10<sup>3</sup>/uL | 39.97 | 2.09–763.23 | 0.014* | Increased hazard |
| Eosinophils (%) | % | 1.47 | 1.19–1.82 | <0.001* | Increased hazard |
| Basophils | 10<sup>3</sup>/uL | 0.70 | 0.09–5.27 | 0.725 | NS |
| Basophils (%) | % | 1.16 | 0.39–3.47 | 0.792 | NS |
| **Erythrogram** |  |  |  |  |  |
| Red blood cell count (RBC) | 10<sup>6</sup>/uL | 0.34 | 0.22–0.52 | <0.001* | Reduced hazard |
| Hemoglobin | g/dL | 0.60 | 0.49–0.74 | <0.001* | Reduced hazard |
| Hematocrit | % | 0.85 | 0.79–0.90 | <0.001* | Reduced hazard |
| Mean corpuscular volume (MCV) | fL | 1.03 | 0.89–1.20 | 0.689 | NS |
| Mean corpuscular hemoglobin (MCH) | pg | 1.25 | 0.80–1.96 | 0.326 | NS |
| Mean corpuscular hemoglobin concentration (MCHC) | g/dL | 1.70 | 0.83–3.47 | 0.144 | NS |
| Red cell distribution width (RDW) | % | 0.66 | 0.39–1.12 | 0.126 | NS |
| Nucleated red blood cells (NRBC, %) | % | 0.57 | 0.15–2.09 | 0.394 | NS |
| **Platelet parameters** |  |  |  |  |  |
| Platelet count | 10<sup>3</sup>/uL | 1.00 | 1.00–1.01 | 0.027* | Increased hazard |
| Mean platelet volume (MPV) | fL | 0.67 | 0.35–1.29 | 0.227 | NS |

Values are from univariable Cox proportional hazards models for continuous predictors of 3-year survival. HRs are reported per 1-unit increase in each parameter. HR, hazard ratio; CI, confidence interval; NS, not statistically significant ( $p \geq 0.05$ ).
\*statistically significant

A 10-marker model provided the strongest overall predictive performance in the present dataset and was carried forward for additional analyses. Among animals with complete data for all 10 continuous predictors (n = 63; deaths within 3 years = 16), the multivariable Cox proportional hazards model including REE, fat percentage, lean mass, fluid, SUN, creatinine, phosphorus, hemoglobin, RBC, and age was significant overall (likelihood ratio χ² = 41.373, df = 10, p < 0.001). The model showed strong discriminatory performance, with a concordance of 0.881 (SE = 0.051), and explained a substantial proportion of variation in survival (R² = 0.481). In the fully adjusted model, with all 10 variables assessed simultaneously in a multivariable analysis, only SUN and RBC remained independently associated with survival. Higher SUN was associated with increased hazard (HR=1.03, 95% CI 1.00–1.05, p=0.023) and higher RBC was associated with reduced hazard (HR=0.09, 95% CI 0.01–0.87, p=0.037). The remaining predictors were not significant after adjustment (all p≥0.129).

### Objective 2: ROC-derived risk thresholds and Kaplan–Meier 3-year survival differences

The 10 markers included in the multivariable Cox model were assessed with ROC analysis to identify the highest performing cut-points (Youden index) for dichotomizing each marker into low versus high-risk categories. Categorization of the variables allowed the construction of a composite risk score (objective 3). Table 5 summarizes the resulting thresholds and AUC values. Several variables that were not significant as continuous predictors demonstrated significant survival discrimination after ROC-based dichotomization, indicating threshold-dependent rather than linear associations with mortality risk. The strongest discrimination was observed for SUN and RBC (both AUC=0.847), followed by hemoglobin (AUC=0.784), creatinine (AUC=0.745), and fluid (AUC=0.735). Using these ROC-derived thresholds, Kaplan–Meier analyses demonstrated significant survival differences (log-rank and complementary tests) for most health markers. Survival differed between low and high-risk groups for REE (log-rank p=0.027), lean mass (p=0.003), fluid (p<0.001), SUN (p<0.001), creatinine (p<0.001), phosphorus (p<0.001), hemoglobin (p<0.001), RBC (p<0.001), and age (p=0.045). Fat percentage showed mixed results and borderline separation (log-rank p=0.064), with Gehan and Tarone–Ware tests reaching p=0.048.

**Table 5.** ROC curves results and thresholds identified as high risk of death.

| Parameter | High Risk Threshold | Youden's Index | AUC |
| --- | --- | --- | --- |
| SUN | $\geq 26$ mg/dL | 0.616 | 0.847 |
| RBC | $\leq 6.69$ ($10^6$/uL) | 0.566 | 0.847 |
| Hemoglobin | $\leq 10.2$ g/dL | 0.547 | 0.784 |
| Creatinine | $\geq 0.5$ mg/dL | 0.531 | 0.745 |
| Fluid | $\geq 3.54$ g | 0.493 | 0.735 |
| Phosphorus | $\geq 4.4$ mg/dL | 0.364 | 0.663 |
| REE | $\leq 26$ kcal/day | 0.321 | 0.657 |
| Lean Mass | $\leq 359.29$ g | 0.389 | 0.644 |
| Fat Percent | $\leq 3.81\%$ | 0.278 | 0.619 |
| Age | $\geq 10$ years | 0.241 | 0.574 |

### Objective 3: Development and evaluation of a composite risk-factor scale

The 10-item composite risk-factor scale included REE, lean mass, fat percentage, fluid, SUN, creatinine, phosphorus, hemoglobin, RBC, and age, and was generated by summing the presence (1) or absence (0) of each ROC-defined high-risk factor (range 0–10). In the analytic sample for the 10-item scale evaluation (n=66; deaths within 3 years=18), the mean (SD) score was 3.3 (2.7). As a continuous predictor, the risk-factor score strongly predicted survival in Cox regression with good discrimination (concordance=0.835), explained approximately 42% of the variance in survival, and estimated that for every risk factor accrued, the risk of death increased 1.75 times (HR=1.75 per 1-point increase, 95% CI 1.43–2.14, p<0.001). Consistent with the time-to-event findings, the score also predicted death status in logistic regression (OR=2.08 per 1-point increase, 95% CI 1.47–2.94, p<0.001; McFadden R²=0.405). In linear regression, higher risk burden was associated with shorter survival time (−2.79 months per 1-point increase, 95% CI −3.54 to −2.04, p<0.001; adjusted R²=0.455).

#### ROC threshold for high-risk classification (scores ≥7) and screening performance

ROC analysis applied to the composite scale identified ≥7 risk factors as the most predictive threshold for classifying individuals as high risk. Kaplan–Meier analysis using this dichotomy showed marked separation (log-rank χ²=62.8, p<0.001). The high-risk group exhibited a mean survival reduction of 19 months (95% CI −24.6 to −13.4, p<0.001; adjusted R²=0.41), in reference to the low-risk group. Estimated survival probabilities further illustrated the gradient. In the high-risk group, survival declined from 63.6% at 12 months, to 18.2% at 24 months, and 0% by 36 months. In contrast, survival in the low-risk group remained at 96.4% at 12 months, 92.7% at 24 months, and 86.7% at 36 months. When evaluated as a screening tool for death within 3 years, the ≥7 threshold produced 11 true positives and 0 false positives, with 7 false negatives and 48 true negatives. This corresponded to 89.4% accuracy, 61.1% sensitivity, 100.0% specificity, 100.0% positive predictive value, and 87.3% negative predictive value.

#### Three-level risk stratification (scores 0–1, 2–6, ≥7)

To improve interpretability across the full score distribution, ROC-based refinement yielded three strata: low (0–1), medium (2–6), and high (7–10). Frequencies by outcome demonstrated complete separation at the extremes: 0% deaths in the low-risk group (18/18 alive), 18.9% deaths in the medium-risk group (7/37), and 100% deaths in the high-risk group (11/11). Three-level Kaplan–Meier analysis confirmed robust differences across groups (overall log-rank χ²=64.5, df=2, p<0.001). Pairwise comparisons showed borderline separation between low and medium risk (log-rank p=0.049) and strong separation for low vs high and medium vs high (both p<0.001) (Figure 1). Table 6 contains a scoring sheet for the 10-item risk-factor scale.

**Figure 1.**
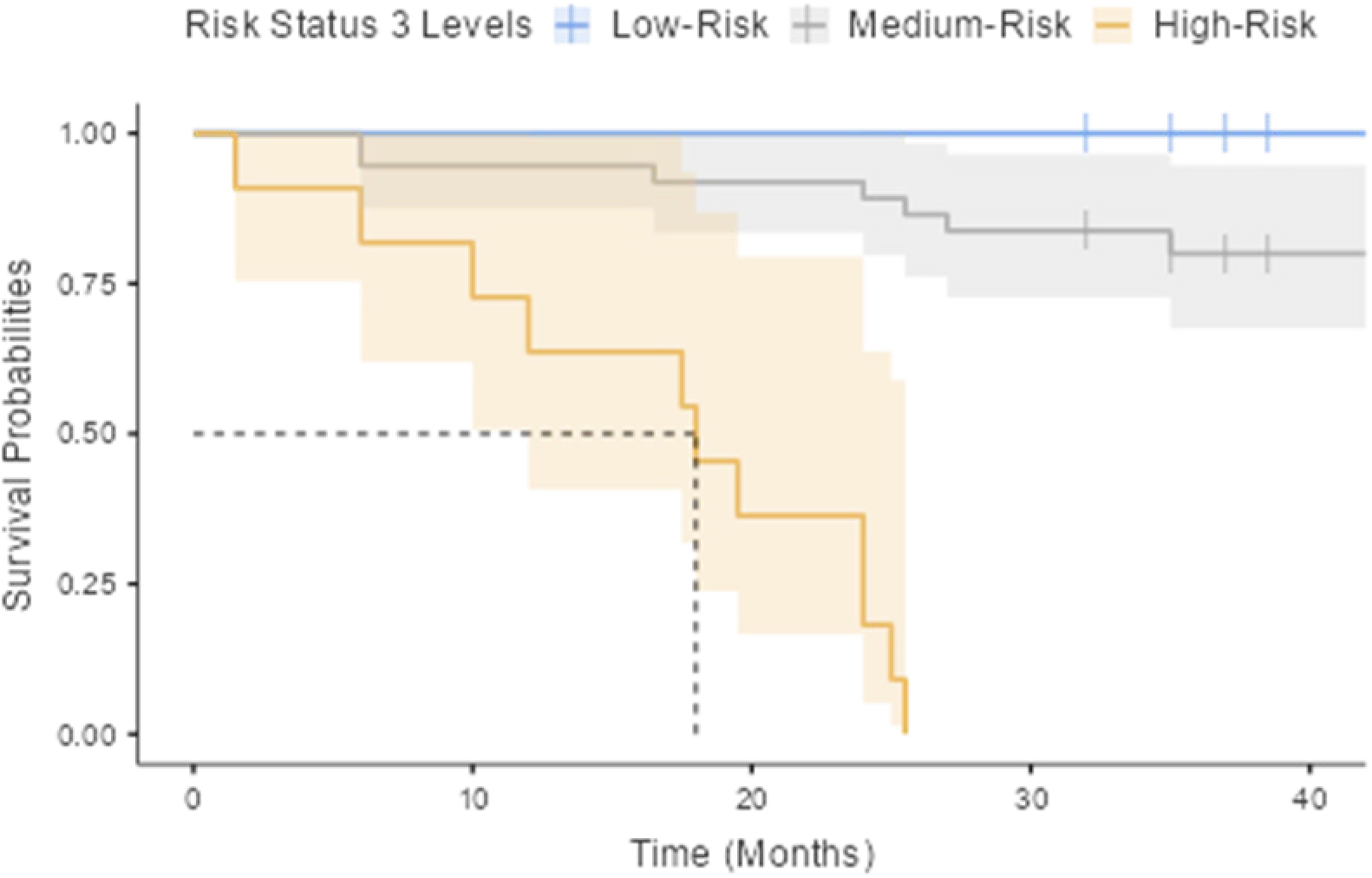
Survival curves for three levels of risk. Low-risk (n=18), medium-risk (n=37), high-risk (n=11).

**Table 6.**
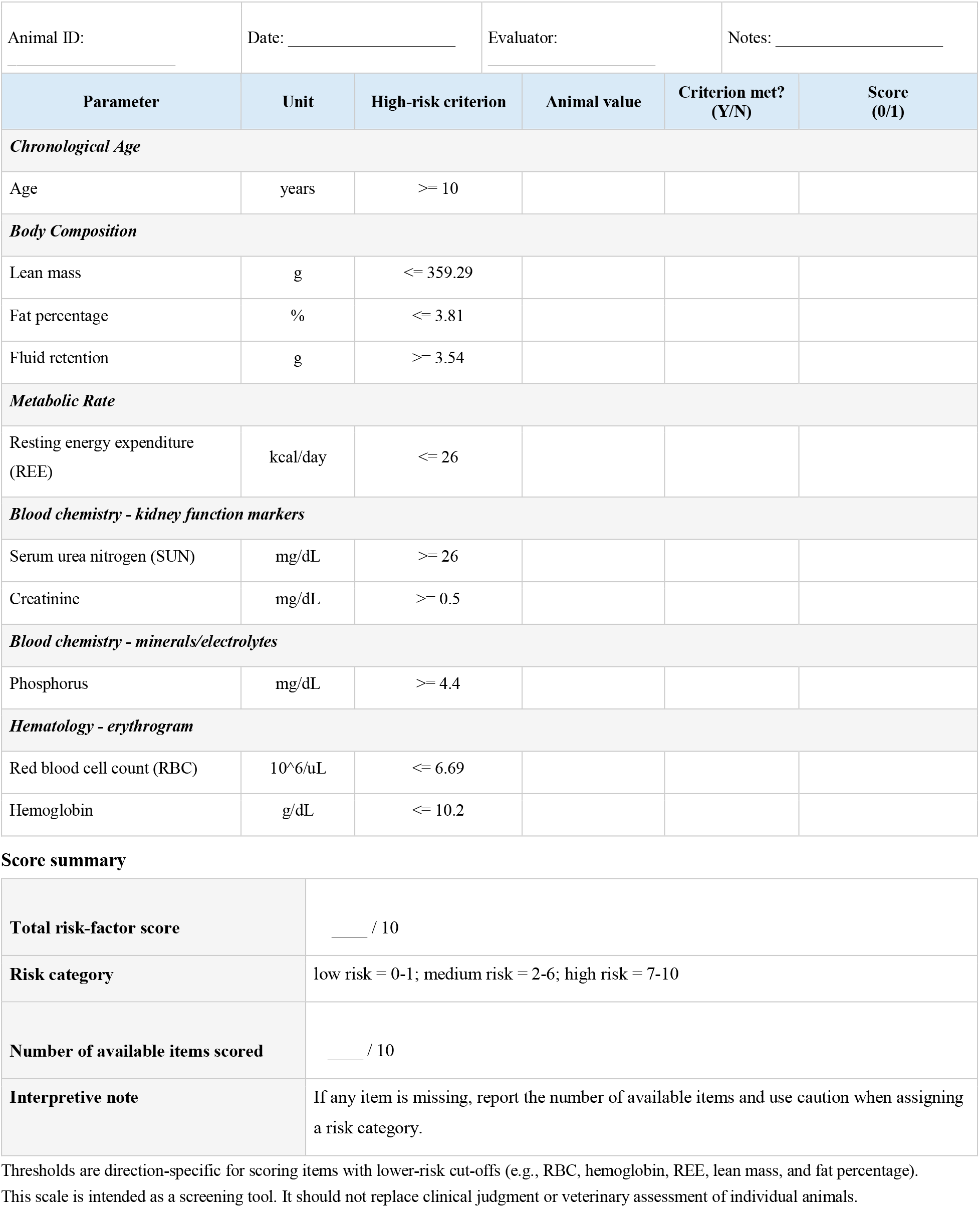
Scoring sheet for the 10-item risk-factor scale for common marmosets. Each parameter is scored as 1 when the high-risk criterion is met and 0 when the criterion is not met. The total score is the sum of the 10 item scores.

|  |  |  |  |  |  |
| --- | --- | --- | --- | --- | --- |
| Animal ID: _____ | Date: _____ | Evaluator: _____ | Notes: _____ |  |  |
| **Parameter** | **Unit** | **High-risk criterion** | **Animal value** | **Criterion met? (Y/N)** | **Score (0/1)** |
| ***Chronological Age*** |  |  |  |  |  |
| Age | years | $\geq 10$ | | | |
| ***Body Composition*** |  |  |  |  |  |
| Lean mass | g | $\leq 359.29$ | | | |
| Fat percentage | % | $\leq 3.81$ | | | |
| Fluid retention | g | $\geq 3.54$ | | | |
| ***Metabolic Rate*** |  |  |  |  |  |
| Resting energy expenditure (REE) | kcal/day | $\leq 26$ | | | |
| ***Blood chemistry - kidney function markers*** |  |  |  |  |  |
| Serum urea nitrogen (SUN) | mg/dL | $\geq 26$ | | | |
| Creatinine | mg/dL | $\geq 0.5$ | | | |
| ***Blood chemistry - minerals/electrolytes*** |  |  |  |  |  |
| Phosphorus | mg/dL | $\geq 4.4$ | | | |
| ***Hematology - erythrogram*** |  |  |  |  |  |
| Red blood cell count (RBC) | $10^6/\mu\text{L}$ | $\leq 6.69$ | | | |
| Hemoglobin | g/dL | $\leq 10.2$ | | | |

**Score summary**
|  |  |
| --- | --- |
| **Total risk-factor score** | ____ / 10 |
| **Risk category** | low risk = 0-1; medium risk = 2-6; high risk = 7-10 |
| **Number of available items scored** | ____ / 10 |
| **Interpretive note** | If any item is missing, report the number of available items and use caution when assigning a risk category. |

Thresholds are direction-specific for scoring items with lower-risk cut-offs (e.g., RBC, hemoglobin, REE, lean mass, and fat percentage). This scale is intended as a screening tool. It should not replace clinical judgment or veterinary assessment of individual animals.

#### Path analysis of physiological domains

A path model (Table 7 and Figure 2) was estimated to summarize directional associations among key predictors across age, body composition (i.e., lean mass, fat%, fluid retention), kidney disease (i.e., SUN, creatinine, fluid retention), anemia (i.e., hemoglobin), and metabolic rate (i.e., REE) domains. The model showed excellent fit (χ²(15)=15.34, p=0.427; CFI=0.998; TLI=0.996; RMSEA=0.019 [95% CI 0.000–0.121]; SRMR=0.065), and fit was markedly better than the baseline independence model (baseline χ²(28)=178.13, p<0.001). Across endogenous variables, explained variance (R²) ranged from 0.094 to 0.506, with the largest R² for kidney and anemia markers (creatinine R²=0.506; hemoglobin R²=0.448) and moderate R² for metabolic rate and body composition/fluid (REE R²=0.350; fluid retention R²=0.220; lean mass R²=0.198; fat% R²=0.197; SUN R²=0.094).

**Figure 2.**
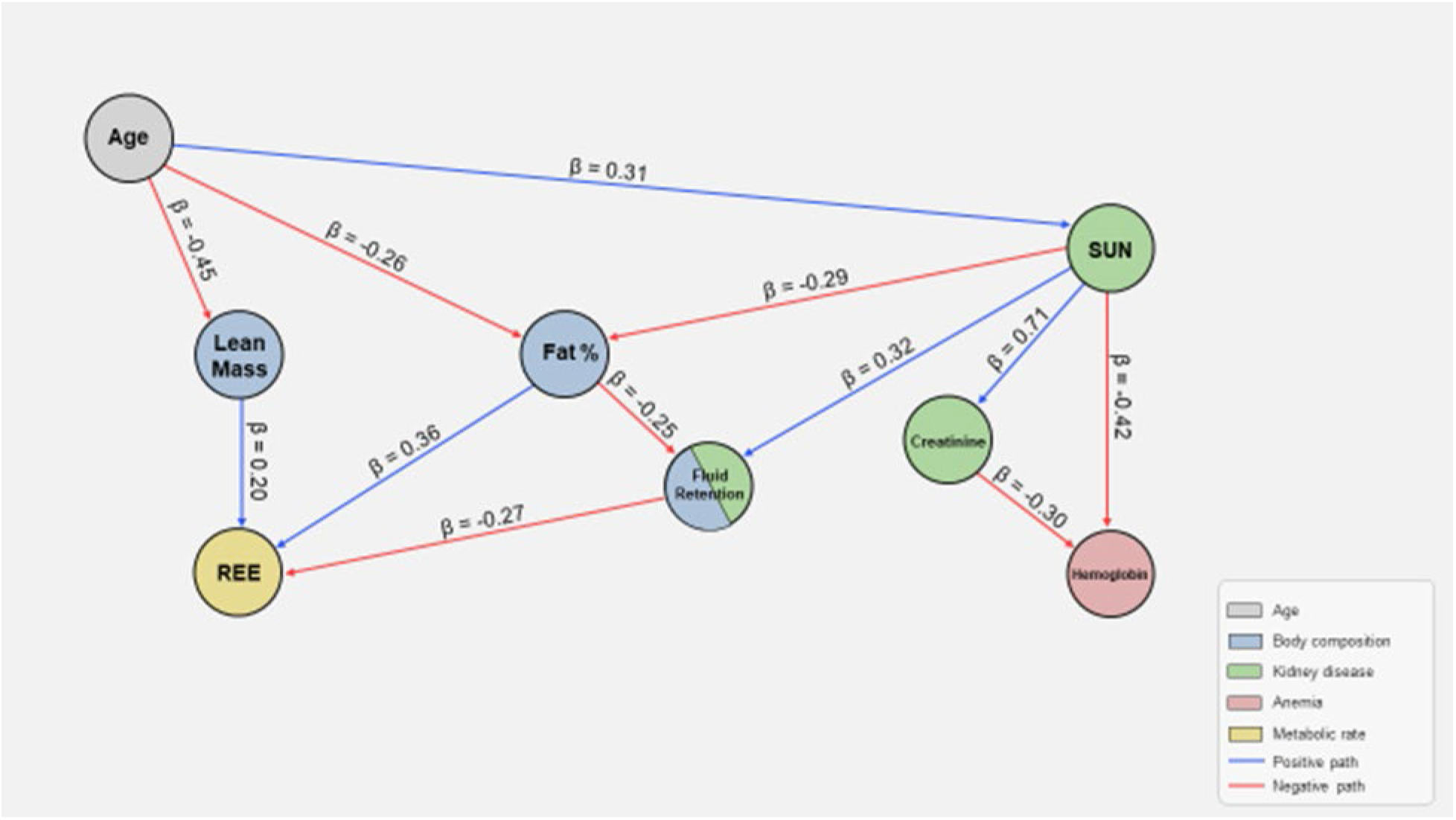
Node diagram for the path analysis model. Standardized path coefficients (β) are shown on arrows. Blue arrows indicate positive associations, and red arrows indicate negative associations. The model includes eight observed variables organized into five color-coded domains: age, body composition, kidney disease, anemia, and metabolic rate. Older age was associated with lower lean mass and fat percentage, and with higher serum urea nitrogen (SUN). Higher SUN was associated with lower fat percentage, greater fluid retention, higher creatinine, and lower hemoglobin. Higher creatinine was also associated with lower hemoglobin. Lower fat percentage was associated with greater fluid retention. In turn, lower lean mass, lower fat percentage, and greater fluid retention were associated with lower resting energy expenditure (REE).

**Table 7.** Path analysis model with age, and parameters of body composition, kidney disease, anemia, and metabolic rate.

| Dependent | Predictor | Estimate | SE | 95% Confidence Intervals | | $\beta$ | z | p |
| --- | --- | --- | --- | --- | --- | --- | --- | --- |
|  |  |  |  | Lower | Upper |  |  |  |
| Lean Mass | Age | -0.445 | 0.1128 | -0.66618 | -0.2239 | -0.445 | -3.94 | <.001* |
| Fat % | Age | -0.260 | 0.1186 | -0.49292 | -0.0279 | -0.260 | -2.20 | 0.028* |
| Fat % | BUN | -0.288 | 0.1186 | -0.52031 | -0.0553 | -0.288 | -2.43 | 0.015* |
| Fluid Retention | BUN | 0.316 | 0.1196 | 0.08167 | 0.5505 | 0.316 | 2.64 | 0.008* |
| Fluid Retention | Fat % | -0.250 | 0.1196 | -0.48425 | -0.0154 | -0.250 | -2.09 | 0.037* |
| BUN | Age | 0.306 | 0.1199 | 0.07114 | 0.5413 | 0.306 | 2.55 | 0.011* |
| Creatinine | BUN | 0.319 | 0.0398 | 0.24138 | 0.3973 | 0.711 | 8.03 | <.001* |
| Hemoglobin | BUN | -0.423 | 0.1319 | -0.68158 | -0.1647 | -0.425 | -3.21 | 0.001* |
| Hemoglobin | Creatinine | -0.660 | 0.2935 | -1.23557 | -0.0852 | -0.297 | -2.25 | 0.024* |
| REE | Lean Mass | 0.198 | 0.0998 | 0.00271 | 0.3941 | 0.203 | 1.99 | 0.047* |
| REE | Fat % | 0.354 | 0.1070 | 0.14441 | 0.5637 | 0.362 | 3.31 | <.001* |
| REE | Fluid Retention | -0.265 | 0.1060 | -0.47237 | -0.0567 | -0.271 | -2.49 | 0.013* |

$\chi^2(15)=15.34$ , $p=0.427$ . Unstandardized path coefficients (Estimate), standardized path coefficients ( $\beta$ ), and significance levels for each parameter in the path analysis model. \*statistically significant

##### Age domain

Age functioned as an upstream factor with direct paths into both body composition and kidney disease. Older age predicted lower lean mass (β=−0.445, p<0.001) and lower fat% (β=−0.260, p=0.028), and independently predicted higher SUN (β=0.306, p=0.011).

##### Body composition domain (lean mass, fat%, fluid retention)

Within body composition, fat% was positioned as a contributor to fluid status. Lower fat% predicted higher fluid retention (β=−0.250, p=0.037). Lean mass did not directly predict fluid retention in the specified model, but contributed downstream to metabolic rate.

##### Kidney disease domain (SUN, creatinine, fluid retention)

Kidney-related markers formed a central substructure in which SUN acted as a key upstream node. Higher SUN predicted higher creatinine (β=0.711, p<0.001) and greater fluid retention (β=0.316, p=0.008). SUN was also associated with lower fat% (β=−0.288, p=0.015), linking kidney disease processes to body composition in the model.

##### Anemia domain (hemoglobin)

Hemoglobin was negatively associated with kidney disease markers. Higher SUN predicted lower hemoglobin (β=−0.425, p=0.001), and creatinine contributed additional independent prediction of lower hemoglobin (β=−0.297, p=0.024). A residual covariance between hemoglobin and REE was not significant (β=0.116, p=0.359), suggesting that their shared variance was adequately captured by the modeled predictors rather than unexplained correlation.

##### Metabolic rate domain (REE)

REE was jointly explained by body composition and fluid retention. Higher REE was associated with greater lean mass (β=0.203, p=0.047) and higher fat% (β=0.362, p<0.001), while greater fluid retention predicted lower REE (β=−0.271, p=0.013).

Overall, the domain-organized path model supported a structure in which age predicted adverse shifts in body composition and increased SUN, and SUN linked kidney disease to downstream correlates of fluid retention and anemia, while REE reflected combined contributions of lean mass, adiposity, and fluid status.

## Discussion

This study aimed to: 1) identify health markers that predict 3-year survival in the common marmoset, 2) translate significant predictors into interpretable risk thresholds, and 3) integrate multi-domain information into a composite risk-factor scale suitable for geroscience research and clinical decision-making in captive colonies. Overall, two general patterns emerged from these analyses. First, while multiple markers spanning metabolic rate, body composition, hematology, and blood chemistry predicted survival in univariable models, renal nitrogen handling and erythrocyte-related measures emerged as the strongest signals. That is, renal disease predominated in the high risk group as reflected by elevations in SUN and chronic anemia. In the fully adjusted Cox model, higher SUN independently increased mortality hazard, whereas higher RBC independently reduced hazard. Second, the aggregate burden of dysregulation operationalized as a 0–10 count of high-risk thresholds showed a strong, graded association with mortality risk and survival time, and identified a small subset of individuals (≥7 risk factors) with extreme mortality risk. These findings align with a geroscience framework in which aging is expressed as multi-system loss of resilience and altered inter-organ communication, rather than isolated dysfunction within a single physiological axis [46].

### Kidney–anemia–fluid axis as a high-salience mortality pathway

A major implication of these data is that kidney-linked processes exhibited high salience in terms of vulnerability in aging marmosets. In humans and other models, reduced kidney function is a convergent endpoint for multiple aging hallmarks such as chronic inflammation, altered intercellular communication, mitochondrial dysfunction, and dysregulated nutrient sensing, which collectively erode physiological reserve and amplify risk across organ systems [23, 47]. The results are consistent with this view. SUN displayed strong discriminatory performance (high AUC) and remained an independent predictor in the multivariable survival model, despite covariation with other domains. In the context of geroscience, SUN can be interpreted as a pragmatic, integrative marker that reflects the combined influence of renal clearance, protein catabolism, fluid volume status, and systemic illness burden. In this sense, decline in kidney function likely suggests a significant transition in physiological aging among marmosets.

The path analysis strengthens the geroscience framing by showing that SUN sits upstream of several downstream liabilities. Higher SUN predicted higher creatinine (i.e., kidney disease domain), greater fluid retention (i.e., body composition and kidney disease domains), and lower hemoglobin (i.e., anemia domain), with creatinine providing additional independent prediction of hemoglobin. This structure is consistent with a renal–hematologic coupling in which declining renal function and/or renal stress contributes to anemia through reduced erythropoietin signaling, inflammatory iron restriction, and reduced red cell survival, representing well-described pathways in aging [24, 48]. Importantly, the marmoset literature supports the biological plausibility of kidney aging as a major contributor to late-life risk. Aged marmosets show renal histopathology (e.g., glomerulosclerosis, interstitial fibrosis, arteriolosclerosis) and functional changes such as increased urinary albumin and protein excretion [12]. Although in the present study we operationalized renal risk via SUN and serum creatinine rather than urinary measures, the convergence between survival predictors and known age-related renal phenotypes supports the interpretation that renal aging is not merely incidental, but a plausible driver of vulnerability.

### Anemia-related signals as markers of resilience

Red blood cell count variability was notable for two reasons. It showed strong discriminatory capacity in ROC curve analysis and remained independently predictive in the multivariable Cox model. In a geroscience framing, this suggests that erythrocyte-related measures may index intrinsic capacity [49] or a broader resilience phenotype, plausibly linked to oxygen delivery capacity, marrow function, inflammatory tone, and kidney–erythropoietic coupling. In older human adults, anemia is associated with frailty, reduced function, and higher mortality risk, as mechanistically chronic inflammation and chronic disease can suppress erythropoiesis and alter iron handling [24, 48]. In marmosets, hematological and biochemical markers show age-related shifts, indicating that age-calibrated “normal” reference ranges may be necessary to distinguish healthy aging from pathological deviation, particularly when interpreting RBC and hemoglobin in older animals [25].

In the present study, the high-risk thresholds identified for SUN (≥26 mg/dL), creatinine (≥0.5 mg/dL), phosphorus (≥4.4 mg/dL), and hemoglobin (≤10.2 g/dL) overlapped with the ranges reported by Hickmott and colleagues [25] for healthy peri-geriatric and geriatric marmosets. However, mean values in those age groups did not exceed these high-risk thresholds. Thus, as healthy marmosets age, these parameters appear to approach, but on average do not exceed, the high-risk thresholds identified here, supporting the need for age-specific reference ranges. A notable exception was RBC, with the high-risk threshold in the present study (≤6.69 × 10^6/µL) exceeding mean RBC values reported in Hickmott, Cervantes [25] for healthy peri-geriatric and geriatric animals. This pattern highlights anemia as a pervasive issue in the sampled population, and suggests that RBC may be a more sensitive anemia-related predictor of mortality risk than hemoglobin, consistent with mechanisms such as chronic inflammation, impaired nutrient absorption, and bone marrow dysfunction.

The path analysis offered a coherent integration of these factors. Kidney markers (i.e., SUN and creatinine) predicted lower hemoglobin, suggesting that the anemia signal is not purely independent but partially embedded within renal and systemic aging processes. Meanwhile, the lack of a significant residual covariance between hemoglobin and REE in the final path model suggests that once renal function and fluid/body composition pathways are specified, additional unexplained coupling between anemia and metabolic rate may be limited. This is conceptually consistent with a geroscience view in which multiple aging hallmarks converge on a smaller number of observable state variables that summarize resilience (e.g., renal clearance, hematologic reserve, fluid homeostasis).

### Body composition, fluid status, and metabolic rate: downstream integration rather than primary drivers

Several body composition and metabolic markers predicted survival in univariable analyses and showed clear Kaplan–Meier separation after dichotomization, notably lean mass and fluid. However, they were attenuated in the fully adjusted Cox model, which is a pattern expected in a multi-domain aging dataset. Body composition and energy metabolism appeared as integrators of upstream disease processes, and their prognostic signal was reduced when more proximal physiological drivers were included. The path analysis is consistent with this interpretation. REE was jointly explained by lean mass and fat% (positive paths) and fluid retention (negative path), implying that metabolic rate is shaped by a combination of tissue mass and fluid volume dysregulation. In practical terms, this suggests REE may be a sensitive gauge for multisystem status, but not a highly specific marker for mechanistic inference when renal and hematologic variables are available.

Two additional points regarding adiposity and fluid status are relevant from a geroscience perspective. First, the borderline/mixed survival separation for fat% (depending on test) may reflect nonlinear and context-dependent effects of adiposity in late life, where both low adiposity (e.g., cachexia/sarcopenia-associated risk) and high adiposity (e.g., metabolic dysregulation) can be detrimental [10, 30]. Second, the negative association between fat% and fluid retention in the path model, after accounting for SUN, underscores that fluid in geriatric animals is not merely a body composition component but may reflect a clinically meaningful state (e.g., edema/volume overload) that sits at the intersection of renal function, inflammation, and nutritional status. In humans, frailty in kidney disease populations is closely intertwined with malnutrition–inflammation phenotypes [50]. These findings and similarities highlight marmosets as a useful translational model for aging and the kidney–anemia–fluid axis.

### Composite risk-factor burden as “biological age” or “frailty” signal

The composite risk-factor scale (Table 6) is best understood as a biomarker-centered deficit count that captures multi-domain physiological dysregulation, using objective measures that are feasible in routine colony practice. Conceptually, it parallels the deficit accumulation approach to frailty, in which mortality risk increases as the number of health deficits increases. This is typically operationalized using indices that aggregate diverse deficits from functional and clinical measures [34]. Despite using only 10 risk factors, the scale behaved in a manner consistent with deficit accumulation theory. Each additional risk factor increased mortality risk and shortened survival time, and higher scores clustered into distinctly different risk strata. This supports the interpretation that, in marmosets, a relatively small set of physiologically salient markers may capture a large fraction of increased vulnerability with aging, especially when those markers span domains repeatedly implicated in aging phenotypes (e.g., kidney, anemia, body composition, and metabolic rate) [6, 9–12, 51].

The ROC-derived threshold of ≥7 risk factors identified an extreme risk (i.e., high-risk) subgroup with outstanding specificity and positive predictive value, suggesting a potential role as a triage flag in geroscience studies (e.g., for enhanced monitoring, exclusion from intervention protocols where imminent death could confound endpoints, or enrollment into palliative/welfare treatment). However, the moderate sensitivity indicates that a meaningful fraction of deaths occurred below this extreme threshold, which is consistent with the heterogeneity of aging trajectories emphasized in geroscience. The three-level stratification (0–1, 2–6, ≥7) addresses this by separating a low-risk group, a heterogeneous intermediate group, and a high-risk group with extreme mortality. This structure suggests the presence of an extended healthspan (i.e., period of general health) with a compressed sickspan (i.e., period of morbidity and disability) in the sampled population; where a small subset is overtly susceptible, while another subset remains resilient and a large middle group expresses graded deficits that may be most responsive to interventions targeting aging biology [46, 52].

### Relevance to the common marmoset as a translational geroscience model

The common marmoset has been increasingly positioned as a powerful translational model for geroscience, as it combines primate physiology with a comparatively short lifespan and measurable age-related morbidity patterns [51]. The findings from this work extend this translational value in two ways. First, they identify a coherent set of routine clinical biomarkers that map onto major aging-related disease axes (i.e., renal decline with anemia, body composition change, altered energetics). Second, they provide an empirically derived, interpretable risk scale that could support longitudinal aging studies by enabling: 1) baseline stratification, 2) covariate adjustment for “physiological age”, and 3) selective enrollment for intervention trials based on specific risk levels. This is particularly relevant given ongoing interest in leveraging marmosets for testing geroprotective interventions and for quantifying aging phenotypes across multiple domains of healthspan [6].

### Clinical utility for captive colonies: pragmatic screening and targeted follow-up

The results have immediate implications for colony management and clinical care. A key advantage of the scale is that it relies on measures already common in veterinary monitoring (i.e., CBC, blood chemistry panel, body composition, and metabolic rate where available). The high specificity of the extreme-risk threshold (≥7) supports its use as a conservative flag to identify animals who may benefit from prompt confirmatory evaluation (e.g., renal assessment, fluid volume evaluation, anemia workup) and increased monitoring frequency. The medium-risk group (2–6) may be the most clinically actionable, as it likely contains animals with early renal compromise, evolving anemia, or emerging dysregulation that could be mitigated by targeted husbandry, dietary optimization, hydration strategies, or early diagnostic testing. Importantly, although we found that chronological age (i.e., age ≥10 years significantly increased risk of death) was not among the strongest predictors of survival in marmosets, published work emphasizes that “healthy aging” reference ranges for hematological biomarkers in geriatric animals may differ from young adult reference intervals [25]. Accordingly, we recommend age-aware clinical interpretation when applying threshold-based tools in the assessment of geriatric marmosets.

### Limitations and future directions

Several limitations should shape interpretation and guide follow-up work. First, the multivariable Cox model included a proportionately large set of predictors relative to the number of deaths, increasing the possibility of overfitting and coefficient instability. This may partly explain why multiple univariable predictors attenuated after adjustment and why SUN and RBC remained as the most robust independent signals. A larger sample with a higher proportion of deaths, in conjunction with penalized Cox approaches, bootstrap shrinkage, or cross-validated model selection might strengthen inference and help identify stable predictor sets for future cohorts. Second, ROC-derived thresholds optimized within the same dataset can be optimistic. Temporal validation (e.g., later cohorts) and/or external validation will be necessary to assess generalizability, to recalibrate cut-points if needed, and determine whether thresholds should be stratified by sex or age. Third, dichotomization improves interpretability but sacrifices information. Future work could compare threshold-based and continuous risk approaches (e.g., restricted cubic splines, generalized additive survival models) to better capture nonlinearity, notably for adiposity and REE.

Lastly, path analysis was used to test hypothesized directional relationships among variables and to provide an integrative summary of the multivariable structure of the data, which can in turn generate new hypotheses. However, path models estimated from observational data should be interpreted as plausible causal structures consistent with the data and model assumptions, not as definitive evidence of causality. In future studies, stronger causal inference will require designs that address confounding and temporal ordering (e.g., randomized controlled experiments when feasible, or well-powered longitudinal/quasi-experimental approaches).

Looking forward, the most valuable extension of this work might be longitudinal analyses. Tracking within-individual trajectories of kidney and anemia markers, fluid status, and body composition could identify early inflection points preceding mortality. This might clarify whether the composite score and changes related to kidney disease and anemia anticipate terminal decline, or if these merely reflect late-stage disease. Integrating additional geroscience markers (e.g., inflammaging, lipidomics, microbiome features, or molecular aging clocks) could connect the clinical biomarker scale to upstream hallmarks and provide a bridge between mechanistic geroscience and practical colony medicine [22, 23, 53].

## Conclusion

In summary, this work supports a geroscience model in which 3-year survival in common marmosets is strongly shaped by multi-system physiological dysregulation, with a prominent kidney–anemia–fluid axis and downstream integration into metabolic rate and body composition. The derived thresholds and composite scale provide an interpretable framework for quantifying vulnerability, stratifying aging cohorts, and supporting translational geroscience studies, while also offering a pragmatic screening tool to enhance clinical monitoring and welfare in captive marmoset colonies [6, 12, 51].

## Acknowledgments

We thank Donna Lane-Colón and the marmoset care team at SNPRC. This investigation used resources that were supported by NIH grant R01AG065546, by the base grant to SNPRC (P51OD011133) from the Office of Research Infrastructure Programs, NIH; and by the Office of the Director, NIH under award number S10OD032443.

## Author contributions

Juan Pablo Arroyo: conceptualization (equal), data curation (lead), formal analysis (lead), investigation (lead), methodology (lead), project administration (supporting), supervision (supporting), validation (lead), visualization (lead), writing – original draft (lead), writing – review and editing (equal).

Aaryn C. Mustoe: conceptualization (equal), formal analysis (equal), investigation (equal), methodology (equal), project administration (supporting), supervision (supporting), validation (equal), visualization (equal), writing – original draft (supporting), writing – review and editing (equal).

Kelly R. Reveles: conceptualization (equal), formal analysis (equal), funding acquisition (equal), methodology (equal), project administration (equal), supervision (equal), visualization (equal), writing – review and editing (equal).

Kathleen M. Brasky: conceptualization (supporting), methodology (equal), project administration (equal), resources (equal), supervision (equal), writing – review and editing (supporting).

Donna Perry: conceptualization (supporting), data curation (equal), formal analysis (supporting), investigation (equal), methodology (supporting), project administration (supporting), resources (equal), supervision (equal), validation (equal), writing – review and editing (equal).

Lidia Cervantes: conceptualization (supporting), formal analysis (supporting), investigation (equal), methodology (supporting), project administration (supporting), supervision (supporting), validation (equal), visualization (equal), writing – review and editing (equal).

Addaline Alvarez: data curation (equal), investigation (equal), methodology (supporting), project administration (equal), supervision (supporting), validation (supporting), writing – review and editing (supporting).

Clarissa Hinojosa: data curation (equal), investigation (equal), methodology (supporting), project administration (equal), supervision (equal), validation (supporting), writing – review and editing (supporting).

Jessica Greig: data curation (equal), investigation (equal), methodology (supporting), project administration (equal), supervision (equal), validation (supporting), writing – review and editing (supporting).

Alexana J. Hickmott: data curation (equal), formal analysis (supporting), investigation (equal), methodology (equal), project administration (supporting), supervision (equal), visualization (supporting), writing – review and editing (supporting).

Benjamin J. Ridenhour: conceptualization (supporting), formal analysis (supporting), funding acquisition (equal), methodology (equal), project administration (equal), supervision (equal), visualization (supporting), writing – review and editing (equal).

Katherine R. Amato: conceptualization (supporting), formal analysis (supporting), funding acquisition (equal), methodology (equal), project administration (equal), supervision (equal), visualization (supporting), writing – review and editing (equal).

Michael L. Power: conceptualization (supporting), data curation (equal), formal analysis (supporting), funding acquisition (equal), methodology (equal), project administration (equal), supervision (equal), validation (equal), visualization (supporting), writing – review and editing (equal).

Corinna N. Ross: conceptualization (lead), data curation (equal), formal analysis (equal), funding acquisition (lead), methodology (equal), project administration (lead), supervision (lead), validation (equal), visualization (equal), writing – original draft (equal), writing – review and editing (lead).

## Competing Interests

The authors declare no competing interests.

## Notes

### Competing Interest Statement

The authors have declared no competing interest.

